# Vagal dopaminergic afferents link interoception to trigeminal pain modulation

**DOI:** 10.64898/2026.03.27.714928

**Authors:** Hyeonwi Son, Deoksoo Han, Tingting Li, John Shannonhouse, Eungyung Kim, Cindy Zhou, Md Sams Sazzad Ali, Noreen Baroya, Man-Kyo Chung, Yu Shin Kim

**Affiliations:** Department of Endodontics, School of Dentistry, University of Alabama at Birmingham; Birmingham, AL, USA; Department of Neurobiology, School of Medicine, University of Alabama at Birmingham; Birmingham, AL, USA; Ilsong Institute of Life Science; Seoul, Republic of Korea; Department of Biomedical Gerontology, Graduate School of Hallym University; Chuncheon, Republic of Korea; Department of Neural and Pain Sciences, School of Dentistry, Program in Neuroscience, Center to Advance Chronic Pain Research, University of Maryland Baltimore; Baltimore, MD, USA

**Author notes:** These authors contributed equally to this work (Hyeonwi Son, Deoksoo Han, Tingting Li).

## Abstract

The vagus nerve conveys interoceptive information, yet how specific vagal sensory afferents regulate pain remains unclear. Here, we tested whether vagus nerve stimulation (VNS) modulates temporomandibular disorder (TMD)–related pain. In a mouse model of TMD, auricular VNS (aVNS) attenuated temporomandibular joint (TMJ) pain behaviors and suppressed sensitization of trigeminal nociceptors. We identified a subset of vagal sensory afferents with dopaminergic features that was sufficient to mediate these effects, as selective activation of these afferents recapitulated the analgesic actions of aVNS. These findings highlight an underappreciated peripheral interoceptive pathway and provide a mechanistic framework for targeted neuromodulation in chronic craniofacial pain.

## Introduction

Interoception shapes our behavior by relaying and integrating internal physiological conditions (*1, 2*). Recent studies have expanded the role of interoception beyond homeostatic monitoring, revealing its contribution to adaptive behavioral decisions. We can sense the absorption of energy-rich nutrients in the gut, experience reward, and subsequently develop preferences that guide future food intake (*3–6*). Similarly, internal signals associated with elevated inflammation can trigger compensatory responses that suppress excessive immune activation (*7*). These interoceptive processes are largely conveyed by the vagus nerve. Accordingly, the vagus nerve has emerged as a central anatomical pathway linking internal physiological states to brain circuits that regulate behavior and physiology.

Research aimed at elucidating the function of the vagus nerve has long focused on vagus nerve stimulation (VNS) as a therapeutic strategy for neurological and psychiatric disorders, including treatment-resistant depression and epilepsy (*8*). These clinical efforts have established VNS as a viable neuromodulation approach, culminating in regulatory approval for specific indications. Motivated by accumulating evidence that vagal signaling can influence nociceptive processing (*9–11*), interest in VNS with respect to pain-related conditions has grown. However, the analgesic effects of VNS have predominantly been interpreted as a consequence of non-selective vagal activation and/or downstream efferent parasympathetic mechanisms (*12, 13*). However, emerging data suggest a potential contribution of discrete sensory afferent subtypes to pain modulation, which needs to be fully defined.

Among craniofacial pain disorders, pain associated with temporomandibular disorder (TMD) is one of the most prevalent and disabling sources of orofacial pain. Patients with TMD commonly exhibit persistent mechanical hypersensitivity and jaw dysfunction, leading to substantial impairment of quality of life (*14–17*). Because speaking, eating, and even facial expressions such as smiling can become painful, TMD substantially interferes with basic daily activities; however, current pharmacological and surgical treatments often provide limited or inconsistent relief, highlighting an urgent need for development of rapidly effective, mechanism-based therapies. VNS has been shown to attenuate trigeminal sensitization and migraine-like pain, and more specifically, auricular VNS (aVNS) activates vagal afferent pathways that engage brainstem circuits involved in descending pain inhibition, stress responses, and autonomic regulation (*12, 18–22*). Therefore, aVNS has emerged as a promising neuromodulatory approach for pain and is now being explored in clinical studies of TMD (*23, 24*). Recent in vivo population-level imaging studies in primary and sensory ganglia, predominantly in dorsal root ganglia, have revealed coordinated and spontaneous activity patterns underlying pathological pain states (*25–28*). These findings suggest the possibility that VNS can directly attenuate TMD-related pain and that distinct vagal afferent populations mediate such effects. We have sought to investigate these possibilities.

Here, we identify a distinct subset of vagal sensory afferents that exhibit dopaminergic features and that mediate VNS-induced analgesia in a TMD animal model. We show that aVNS significantly attenuates temporomandibular joint (TMJ) pain behaviors associated with TMD. Importantly, selective activation of dopaminergic-featured vagal afferents innervating the TMJ region is sufficient to suppress pain behaviors and sensitization of trigeminal nociceptors. Together, our findings show that engagement of a specific vagal sensory afferent population is sufficient to modulate trigeminal nociceptors producing analgesia in TMD. These findings provide a mechanistic basis for interoceptive-based neuromodulatory therapies in chronic craniofacial pain.

## Results

### Activation of vagal afferents attenuates FMO-induced pain behaviors

To determine the effects of vagus nerve activation on TMJ pain, we performed aVNS in an established TMD animal model induced by forced mouth opening (FMO) (*29, 30*). We previously established stimulation parameters under which aVNS activates neurons in the nucleus tract solitarius (NTS) (*31*) but not in the trigeminal nucleus caudalis. We modified the method of electrical stimulation by using a concentric electrode attached to a conductive pad, which was placed on the concha of the ear (Fig. 1, A to C). We tested three frequencies of stimulation for analgesic effects on TMJ pain by von Frey assay after 2 day FMO (Fig. 1, D and E). After von Frey measurement, we delivered aVNS. Daily stimuli at 15 Hz did not produce analgesia. Stimulation at 5 Hz produced delayed analgesic effects, which became evident after two consecutive days of daily stimuli. A 2 Hz stimulation produced robust analgesic effects one day after the stimulation (Fig. 1E). These results showed that mechanical thresholds remained elevated the day after aVNS stimulation at 2 Hz or 5 Hz. Notably, the effect emerged the day after stimulation, indicating that aVNS produces analgesia lasting at least 24 h. Even after we stopped administering aVNS following four days of daily aVNS stimulation, EF50 was maintained at an elevated level, especially for aVNS stimulation at 2 Hz. Therefore, we used 2 Hz for the rest of the study (Fig. 1).

**Fig. 1.**
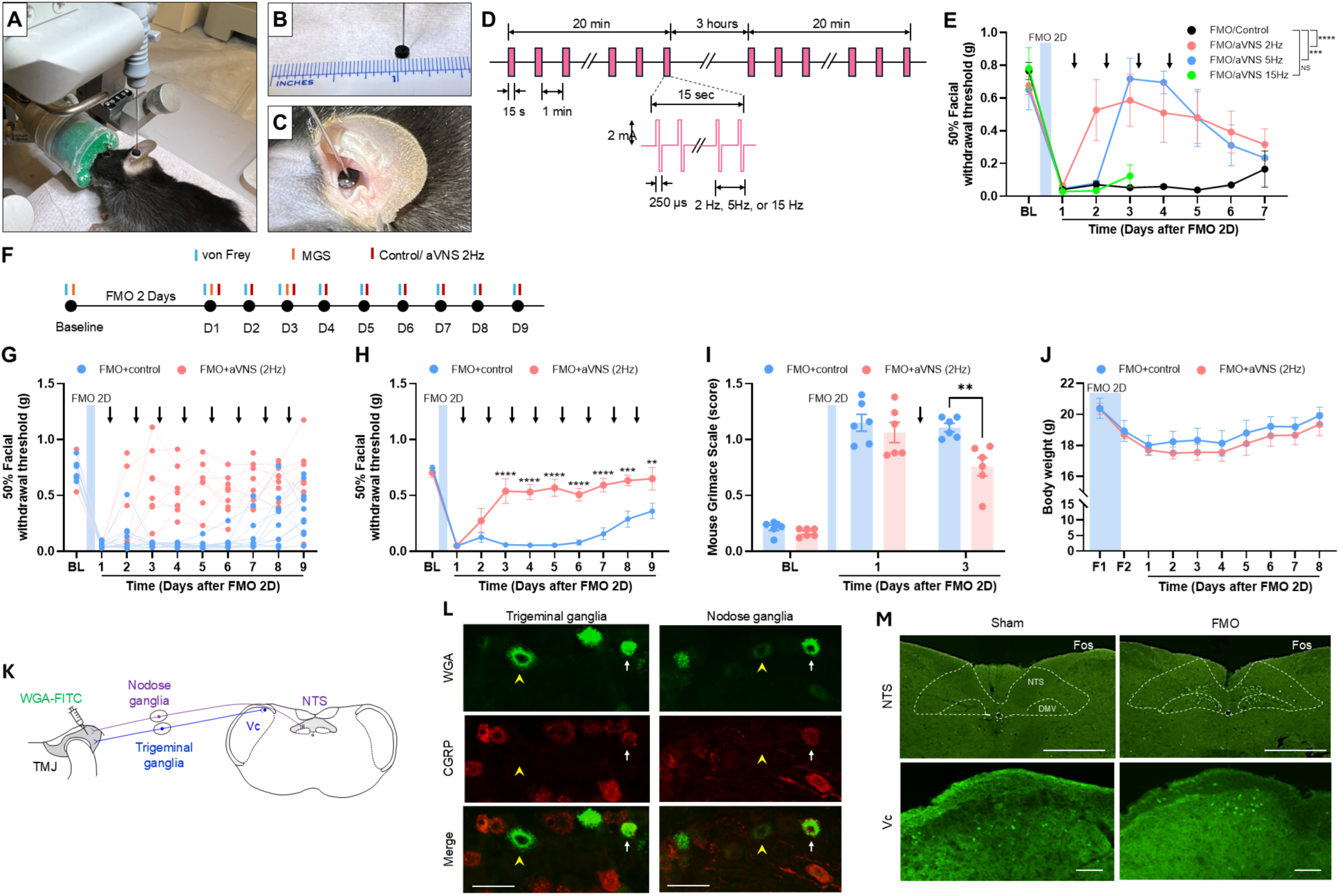
The effects of auricular vagus nerve stimulation (aVNS) on mechanical hyperalgesia, spontaneous pain after forced mouth opening (FMO). (**A**) A mouse under isoflurane anesthesia with the stimulation electrode attached to the ear. (**B**) The dimension of the electrical pad attached to the tip of the concentric electrode. (**C**) The placement of the electrical pad on the conchae of the ear. (**D**) The parameter for the auricular vagus nerve stimulation (aVNS). The daily aVNS is composed of two 20 min sessions at a 3 hour interval. Each 20 min session is composed of electrical stimulation bursts (15 sec duration) at one minute intervals. Each burst is composed of biphasic 2 mA pulses (250 μs duration) at 2, 5, or 15 Hz. (**E**) Von Frey assay was performed in mice following forced mouth opening for 2 days (2 day FMO). The aVNS was delivered at three different frequencies as indicated. aVNS was delivered daily as indicated by arrows. N=4 control animals, 4 treated @ 2 Hz, 3 @ 5 Hz, and 3 @ 15Hz. ***p<0.001; ****p<0.0001; NS, not significant (Sidak’s multiple comparisons of group means followed by two-way ANOVA). (**F**) Time course for testing the effects of aVNS on FMO-induced hyperalgesia. aVNS was applied after performing behavioral assays on each day. (**G, H**) Mechanical threshold (EF50) assessed on the skin overlying left temporomandibular joint. **p<0.01; ***p<0.001; ****p<0.0001 (Sidak post-hoc assay following two-way ANOVA). N=10 controls and 9 aVNS. The arrows indicate aVNS administration. (**I**) Mouse grimace scale (MGS). **p<0.005 (Sidak post-hoc assay following two-way ANOVA). N=6 mice per group. (**J**) Body weight of mice. N=10 controls; 9 aVNS. (**K**) Diagram of dual innervation of TMJ by somatosensory afferents (trigeminal ganglia neurons) and vagal afferents (vagal ganglia neurons). Somatosensory afferents are connected with trigeminal nucleus complex including trigeminal nucleus caudalis (Vc), whereas vagal afferents are connected with nucleus tract solitarius (NTS). Wheat germ agglutinin conjugated with Alexa Fluor 488 (WGA-488) was injected into TMJ to retrogradely label afferents. (**L**) Trigeminal ganglia (left) and vagal ganglia (right) neurons labeled by WGA (top), calcitonin gene-related peptide (CGRP; middle), or merged (bottom). Arrows indicate WGA-CGRP co-expressing neurons; arrowheads indicate neurons labeled by WGA but not expressing CGRP. Scale bar, 50 µm. (**M**) The mice were euthanized two hours after second day of FMO or sham. NTS (upper) or Vc (bottom) regions were immunolabeled using anti-Fos antibody. Dotted lines represent demarcation of nucleus tract solitarius (NTS) and dorsal motor nucleus of the vagus (DMV). Scale bar, 500 µm in NTS and 100 µm in Vc.

We assessed mechanical sensitivity using von Frey and spontaneous pain using mouse grimace scale (MGS) assays (Fig. 1F). FMO induced mechanical hyperalgesia after 1 day (Fig. 1, G and H). After von Frey measurement, we delivered aVNS. On the day following the first aVNS treatment (day 2), some mice (3/9) in the aVNS group showed elevated EF50, but this elevation did not reach statistical significance. On day 3, most mice that received aVNS (7/9) exhibited significantly increased EF50 compared with controls (Fig. 1, G and H). Between day 3 and day 9, EF50 was maintained at similarly high levels in the FMO+aVNS group, whereas the FMO+Control group showed a return of EF50 toward baseline from day 7 to day 9.

We also determined non-evoked pain-like behaviors by assessing MGS. One day after FMO, MGS scores increased and remained elevated in the FMO+Control group through day 3. In contrast, the FMO+aVNS group showed significantly decreased MGS scores on day 3 compared to the FMO+Control group (Fig. 1I). These results showed that aVNS treatment attenuated mechanical hyperalgesia and spontaneous pain-like behaviors following FMO, supporting the anti-nociceptive roles of the activation of vagal ganglia neurons.

Two day FMO resulted in substantially reduced body weight, which gradually recovered toward baseline over a week (Fig. 1J). Body weights of the control and the aVNS groups were not significantly different, supporting the absence of overt adverse effects of aVNS on general health.

We next wanted to determine how the vagal afferent activation influences trigeminal ganglia afferents. Developmental and anatomical studies indicate that the external ear receives somatosensory innervation from multiple cranial nerves, including the auricular branch of the vagus nerve and branches of the trigeminal nerve (*22, 32*). This mixed craniofacial innervation suggests that vagal and trigeminal afferents may exhibit peripheral proximity and potential convergence within adjacent tissues. Based on this rationale, we asked whether vagal afferents are present in the TMJ and whether they are anatomically positioned to interact with trigeminal sensory terminals. We determined the anatomical basis of this interaction in mice by simultaneous labeling of trigeminal ganglia (TG) and nodose ganglion (NG) afferents projecting to the TMJ (Fig. 1K). The injection of wheat germ agglutinin (WGA)-Alexa488, a tracer commonly used for retrograde labeling, into the TMJ resulted in labeling of TG neurons as previously reported (Fig. 1L). In the same mice, we found labeling of NG neurons as well. A proportion of WGA-labeled TMJ-projecting NG afferents were peptidergic afferents co-localizing with calcitonin gene-related peptide (CGRP), as seen in the TG. To further assess the possibility of dual projection of somatosensory and vagal afferents in mouse TMJ, we performed two day FMO and examined Fos expression in the NTS and the trigeminal nucleus caudalis (Vc) (Fig. 1M). Compared to sham, FMO-induced TMJ injury produced robust expression of Fos in the NTS, especially in the medial NTS, and dorsal motor nucleus of the vagus (DMV). These results suggest that vagal afferents project directly to mouse TMJ and suggest that there is peripheral proximity between vagal afferent and trigeminal afferent pathways.

### Dopaminergic vagal afferent activation suppresses TMD-related pain

The vagus nerve contains a diverse set of molecularly defined sensory neuron subpopulations (*33, 34*). Given that aVNS primarily engages vagal afferent pathways22, it is likely that one or more of these subpopulations mediate the analgesic effects of aVNS. Therefore, we sought to identify the afferent subtypes responsible for pain modulation by selectively activating genetically defined vagal sensory neurons in the NG. To accomplish this, we unilaterally injected AAV-DIO-hM3Dq or AAV-DIO-mCherry into the right NG of TH-Cre, TrkC-Cre, MrgD-Cre, CGRP-Cre, and wild-type (WT) mice four weeks prior to the FMO procedure (Fig. 2A). Robust mCherry expression restricted to the ipsilateral NG following unilateral AAV injection confirmed effective and spatially confined viral transduction of vagal sensory neurons (Fig. 2B). To selectively activate Cre-dependent hM3Dq-expressing neurons, we administered Designer Receptors Exclusively Activated by Designer Drugs (DREADD) agonist compound 21 (C21; 0.5 mg/kg) once daily for two consecutive days beginning 24 h after the final FMO session, via intraperitoneal (i.p.) injection in all cre-driver mouse lines except dopamine transporter DAT-cre mice, which received intra-TMJ injection (Fig. 2A). C21 administration produced distinct analgesic effects across Cre lines: TH-Cre mice exhibited significant restoration of facial mechanical thresholds, whereas TrkC-Cre, MrgD-Cre, CGRP-Cre, and WT mice showed no detectable changes (Fig. 2C). These results indicate that the specific afferent subtypes of vagal sensory neurons responsible for pain modulation in the NG express tyrosine hydroxylase (TH).

**Fig. 2.**
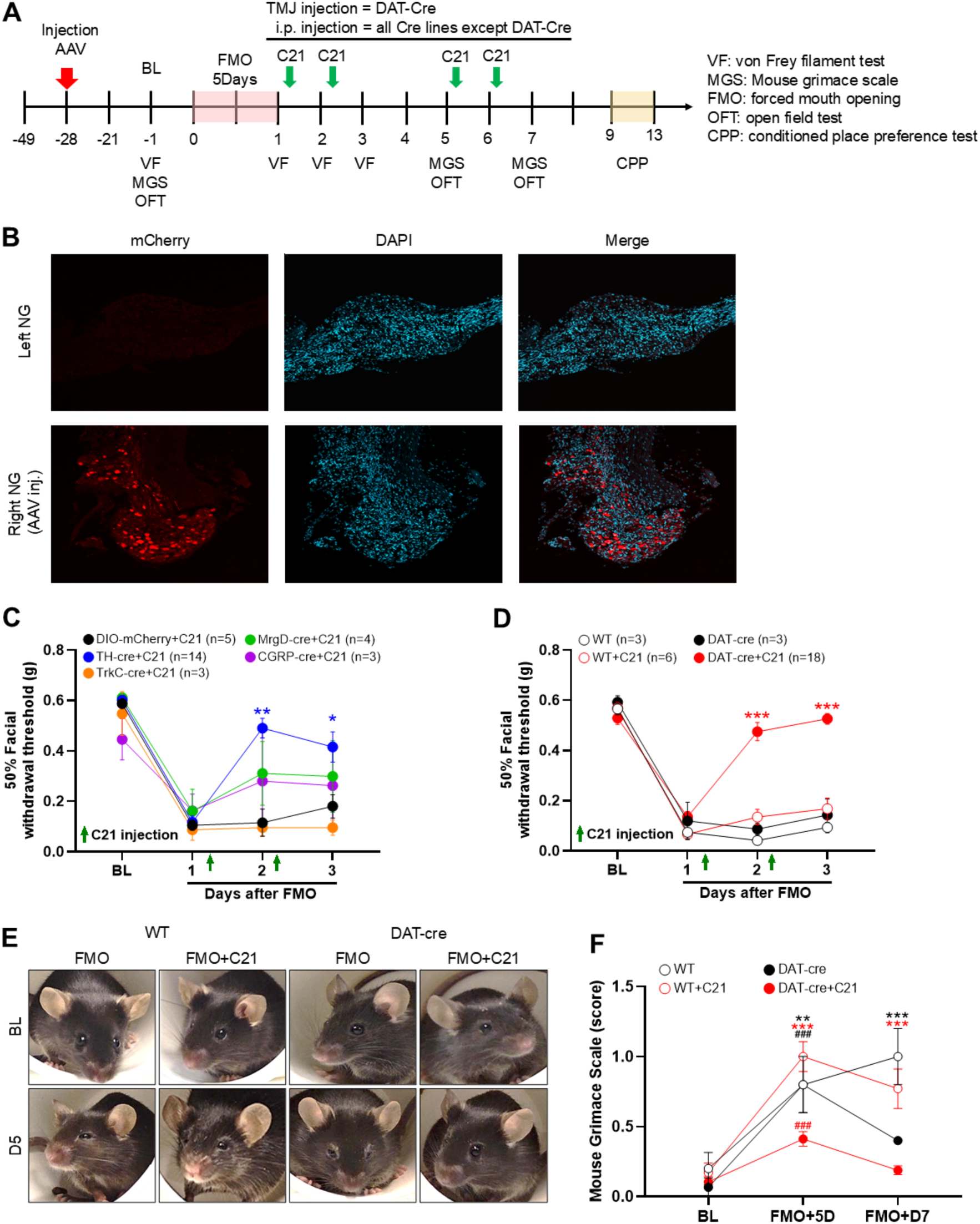
Chemogenetic activation of dopaminergic vagal afferents alleviates mechanical hypersensitivity and spontaneous pain after forced mouth opening (FMO). (**A**) Experimental timeline. AAV vectors were injected into the nodose ganglion (NG) 28 days before the forced mouth-opening (FMO) procedure. FMO was performed for 3 h per day across five consecutive days. Facial mechanical thresholds (von Frey), grimace scores (MGS), and open-field activity were assessed at baseline and multiple time points after FMO. C21 was administered intraperitoneally once daily for 2 days beginning on day 1 post-FMO, and conditioned place-preference (CPP) testing was conducted between days 9–13 post-FMO. (**B**) Representative confocal fluorescence images of NG showing mCherry reporter expression following viral infection, with nuclei counterstained with DAPI. (**C**) Facial withdrawal thresholds across different Cre lines and wild-type after FMO. C21 was administered intraperitoneally after von Frey testing once daily on day 1 and 2 after FMO. Cre-driver mice received nodose ganglion injection of AAV-DIO-hM3Dq and control mice received AAV-DIO-mCherry, both received intraperitoneal C21 after injection. (**D**) Facial withdrawal thresholds in DAT-Cre mice or wild type mice with or without intra-TMJ injection of C21. DAT-cre mice received nodose ganglion injection of AAV-DIO-hM3Dq, followed by intra-TMJ injection of C21 or vehicle. Data are presented as mean ± s.e.m.: *p < 0.05, **p < 0.01, ***p < 0.001 (two-way ANOVA followed by Tukey’s multiple-comparisons test). (**E**) Representative facial images of WT and DAT-Cre mice before and after FMO with or without C21 treatment. (**F**) Quantification of spontaneous pain using the Mouse Grimace Scale (MGS). DAT-Cre + C21 mice exhibited significantly reduced MGS scores on day 5 and day 7 post-FMO compared to control groups. Data are mean ± s.e.m.; **p < 0.01, ***p < 0.001, ###p < 0.001 (two-way ANOVA followed by Tukey’s multiple-comparisons test).

A subset of NG neurons expressing TH and dopa decarboxylase (DDC) are known to comprise a distinct cluster within a transcriptionally defined sensory neuron population33. This suggests that the identified TH-expressing vagal afferents possess dopaminergic molecular features. Given the anatomical and functional proximity between vagal afferents and trigeminal afferents in the TMJ as described in Figure 1, we reasoned that dopaminergic vagal afferents innervating the TMJ could directly contribute to the modulation of trigeminal nociceptive signaling. To directly activate dopaminergic afferents at the vagal nerve terminals, we selectively targeted dopamine transporter (DAT)-expressing vagal afferents. To this end, hM3Dq was expressed in NG neurons via AAV delivery in DAT-Cre mice, and vagal afferent activity was selectively engaged by local administration of C21 into the TMJ, thereby enabling regionally confined activation of a defined vagal sensory afferent population. Consistent with results observed in TH-Cre mice, C21-induced chemogenetic activation significantly attenuated mechanical hypersensitivity and reduced MGS scores in the FMO model (Fig. 2, D and F). Taken together, these findings indicate that activation of dopaminergic vagal afferents produces analgesic effects, suggesting that this population contributes to aVNS-related pain relief.

### Activation of dopaminergic vagal afferents from the orofacial region elicits positive motivational valence

To determine whether activation of dopaminergic vagal afferents alters affective motivational states associated with pain relief, we employed a conditioned place preference (CPP) procedure in which mice learned to associate an environmental context with vagal activation. During conditioning, the striped chamber was paired with C21 (0.5 mg/kg), administered intraperitoneally in all Cre lines except DAT-Cre mice, which instead received intra-TMJ injection, whereas the plain chamber was paired with saline, with the two sessions separated by 6 h (Fig. 3A). On the test day, DAT-Cre mice displayed a robust preference for the C21-paired chamber, reflected by a significant positive ΔCPP score, indicating positive motivational valence associated with dopaminergic vagal activation (Fig. 3, D and E). In contrast, TH-Cre mice exhibited a modest but significant negative ΔCPP score, suggesting an opposing affective valuation. TrkC-Cre, MrgD-Cre, CGRP-Cre, and WT mice showed no detectable change in chamber preference following C21 conditioning (Fig. 3, B and C). Importantly, this CPP effect was not attributable to generalized changes in locomotor or exploratory behavior, as chemogenetic activation of vagal afferents did not produce robust or consistent alterations in open field performance across Cre-driver lines (fig. S1). Collectively, these results identify pronounced cell-type specificity in the affective consequences of vagal afferent activation and indicate that dopaminergic vagal afferents uniquely encode positive motivational valuation, consistent with relief-associated affective modulation rather than nonspecific changes in activity.

**Fig. 3.**
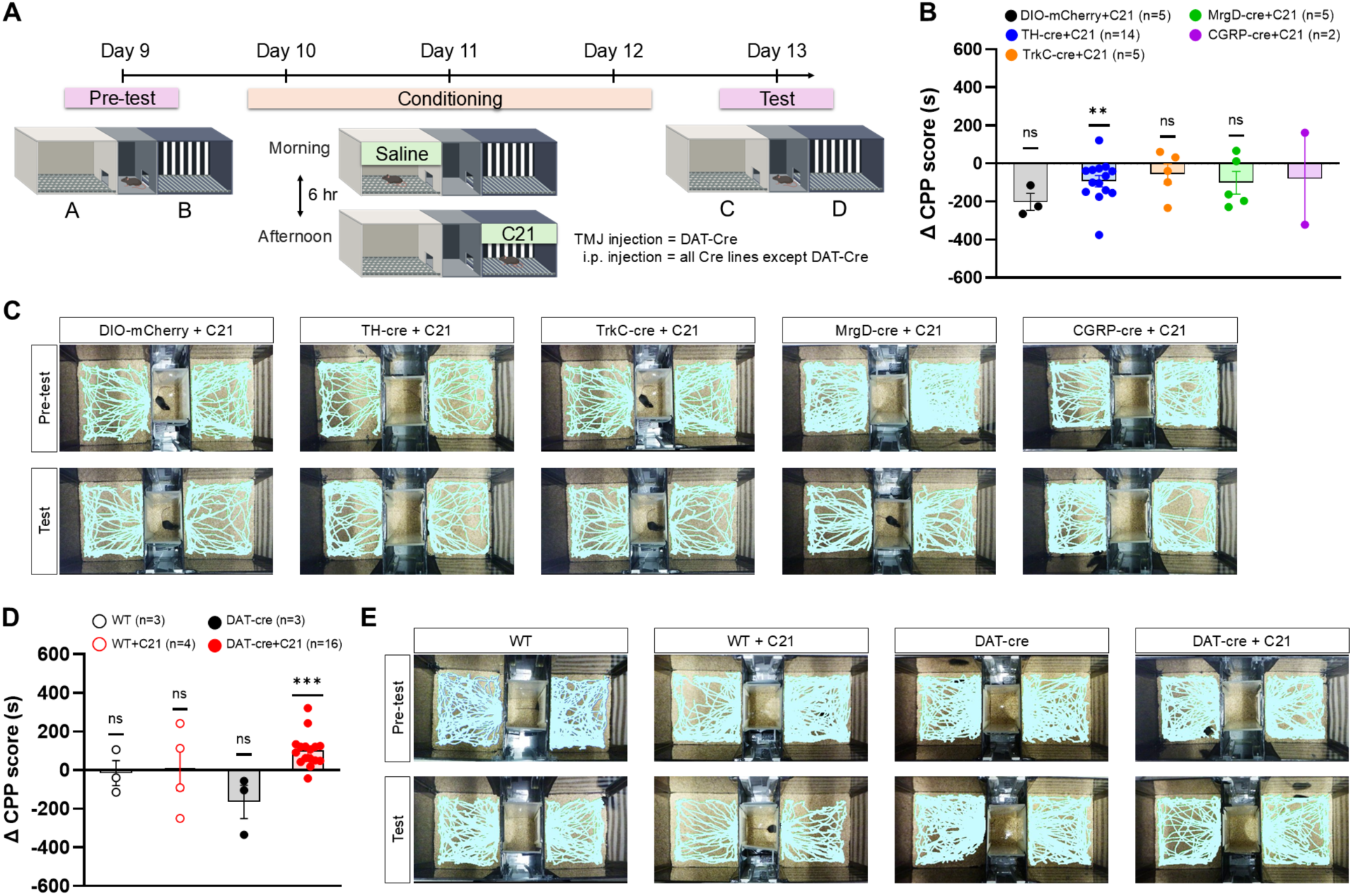
Chemogenetic activation of dopaminergic vagal afferents produces conditioned place preference (CPP). (**A**) CPP experimental timeline. On day 9 (pre-test), mice freely explored both chambers. Days 10-12: Morning saline and afternoon C21 conditioning sessions (6 h apart). Day 13: Test phase with free access to both chambers. (**B**) CPP scores (Δ time in C21-paired chamber) in different Cre lines. (**C**) Representative tracking traces of mice during the pre-test and test sessions in the CPP chamber. (**D**) Comparison of WT and DAT-Cre mice with or without C21 treatment to determine if the CPP effect was specific to DAT-Cre mice. Data are median ± IQR; **p < 0.01, ***p < 0.001 (Wilcoxon signed-rank test vs 0 and Kruskal–Wallis with Dunn’s post hoc). (**E**) Representative tracking traces illustrating changes in chamber preference from the pre-test to the test session in the CPP chamber.

### Dopaminergic vagal activation suppresses spontaneous and evoked trigeminal activity

Our behavioral analyses—including mechanical threshold, MGS, OFT, and CPP—suggest that dopaminergic vagal activation exerts analgesic effects. The anatomical proximity led us to examine whether dopaminergic vagal activation suppresses peripheral sensitization at the level of the TG. FMO markedly increased spontaneous and stimulus-evoked TG activation (*29, 30, 35*). In vivo calcium imaging of intact TG neurons was conducted after completion of the FMO procedure and C21 injection, as outlined in the experimental timeline (Fig. 4A). Consistently, in vivo calcium imaging of intact TG neurons in DAT-Cre/Pirt-GCaMP3 mice with AAV-DIO-hM3Dq injection in NG revealed extensive spontaneous neuronal hyperactivity following FMO-induced TMJ injury (Fig. 4B). Administration of C21 directly into TMJ markedly reduced the number of spontaneously active TG neurons (Fig. 4, C and D, and movie S1 and S2).

**Fig. 4.**
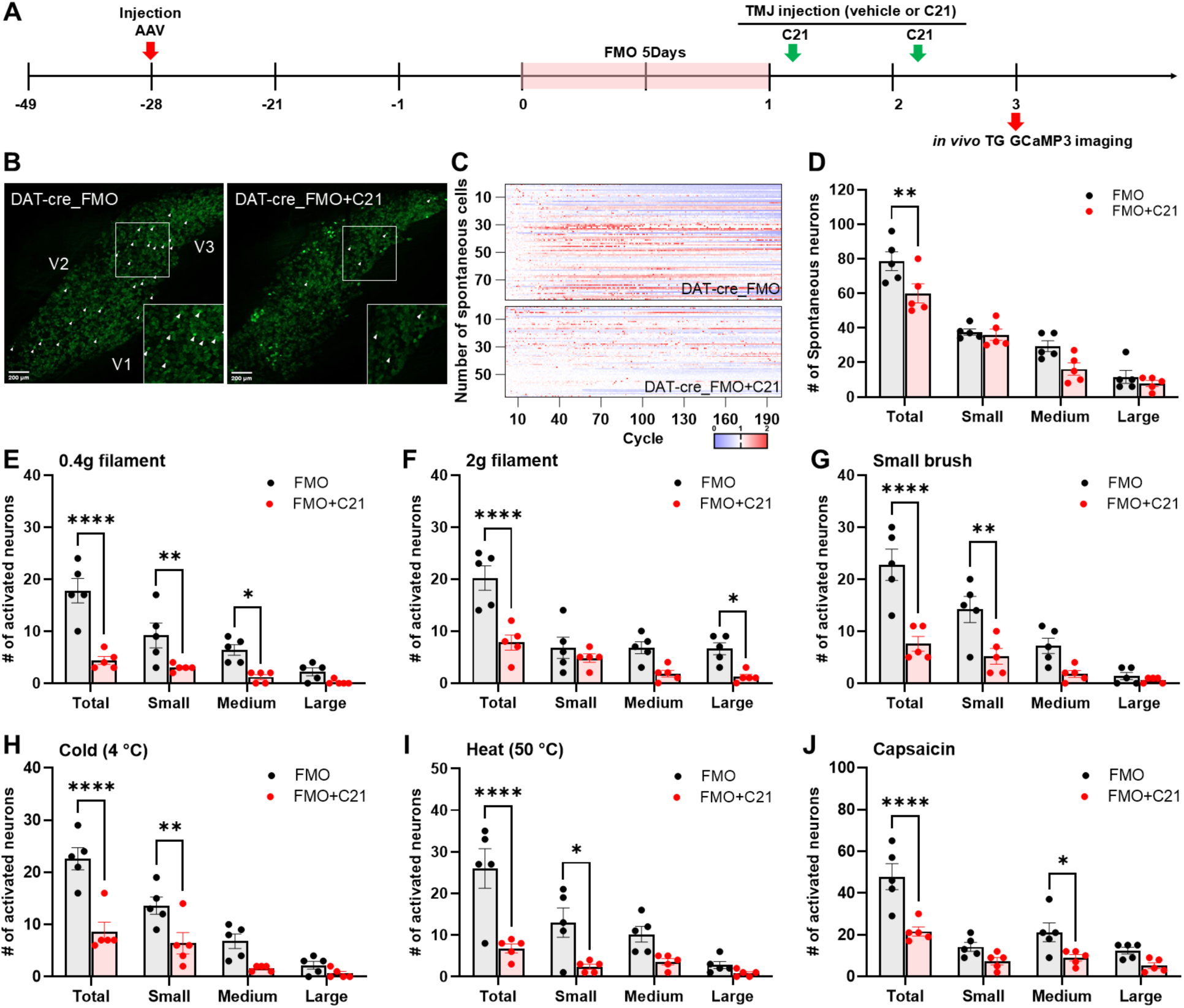
Chemogenetic activation of dopaminergic vagal afferents suppresses spontaneous and stimulus-evoked TG neuronal activity after FMO. (**A**) Schematic illustration of the experimental timeline for in vivo Pirt-GCaMP3 Ca2+ imaging. (**B**) Representative calcium activity maps of trigeminal ganglion (TG) neurons expressing GCaMP3 in DAT-Cre; GCaMP3 mice after FMO with or without C21 treatment. V1–V3 indicate the ophthalmic, maxillary, and mandibular branches of the TG. White arrowheads denote spontaneously active neurons. (**C**) Heat maps showing calcium activity of individual TG neurons over time. (**D**) Quantification of the total number of spontaneously active neurons grouped by soma size (small, medium, large). (**E** to **J**) Quantification of TG neurons activated by different V3 stimuli: (E) 0.4 g filament, (F) 2 g filament, (G) small brush, (H) cold (4°C), (I) heat (50°C), and (J) capsaicin. Data are presented as mean ± s.e.m. (n = 5 mice per group). *p < 0.05, **p < 0.01, ****p < 0.0001 (Two-way ANOVA followed by Tukey’s multiple-comparisons test).

In addition to suppressing spontaneous activity, dopaminergic vagal activation significantly attenuated stimulus-evoked TG responses across multiple sensory modalities. The number of TG neurons responding to mechanical stimulation (0.4 g and 2 g von Frey filaments and small brush) (Fig. 4, E to G, and movie S3 to S8), thermal stimulation (4°C cold and 50°C heat) (Fig. 4, H and I, and movie S9 and S10), and chemical stimulation (capsaicin) (Fig. 4J) were substantially reduced following C21 treatment (Fig. 4, E to J, and movie S11 and S12). Representative superimposed Ca²⁺ transient traces and corresponding area under the curve (AUC) analyses for each stimulus condition are shown. (fig. S2). Collectively, these findings demonstrate that activation of dopaminergic vagal afferents dampens both spontaneous and evoked neuronal hyperactivity within the TG, thereby attenuating peripheral nociceptive drive following FMO-induced TMJ injury.

## Discussion

Vagus nerve stimulation has long been recognized as an effective neuromodulatory approach for pain and affective disorders, although the cellular substrates and sensory mechanisms underlying the therapeutic effects have remained undefined. Our findings suggest that the analgesic effects of vagus nerve stimulation can be mediated by discrete populations of vagal sensory afferents, highlighting a previously underappreciated role of vagal interoceptive signaling in the modulation of craniofacial pain.

The vagus nerve serves as a major conduit for interoceptive signals from peripheral organs to the brainstem. Recent functional studies have shown that genetically defined subsets of vagal sensory neurons can engage arousal-related neuromodulatory systems, shape affective valence, and influence nociceptive processing. Transcriptional profiling of the NG by Kupari and colleagues identified 18 molecularly distinct vagal sensory neuron types, including a notable subtype characterized by the expression of TH and DDC, indicating a dopaminergic molecular signature (*33*). Peripheral neurons expressing TH have been conventionally interpreted as catecholaminergic efferent neurons associated with sympathetic function (*36*). However, some findings indicate that TH-expressing vagal neurons lack the full enzymatic machinery required for classical catecholamine synthesis, distinguishing them from canonical sympathetic efferents (*37*), suggesting that TH-positive NG neurons represent a specialized afferent sensory population, rather than an autonomic motor or efferent population. Consistent with this view, our data indicate that the TH-expressing neuron subgroup participates in an afferent sensory circuit capable of modulating trigeminal nociceptive processing. Rather than acting through direct motor or autonomic efferents, these neurons are likely to convey internal physiological or homeostatic information to central circuits that influence pain sensitivity. This interpretation identifies a cellular basis for how specific vagal afferent populations can modulate pain processing in craniofacial circuits.

Importantly, our findings further reveal functional specialization within TH-expressing vagal afferents. Transcriptional and functional studies have shown that TH+ vagal sensory neurons are molecularly heterogeneous and do not uniformly express the full complement of enzymes and transporters required for functional dopamine signaling. In this context, the observation that chemogenetic activation of TH-Cre neurons attenuated mechanical hypersensitivity but failed to suppress spontaneous pain-like behaviors suggests that this broad TH+ population primarily regulates stimulus-evoked, sensory-discriminative aspects of nociception (*38*).

By contrast, DAT-expressing vagal afferents constitute a more restricted subpopulation with enhanced capacity for dopamine handling and neuro-modulatory signaling. Selective activation of DAT⁺ vagal afferents robustly suppressed both evoked trigeminal responses and spontaneous pain-related behaviors, including affective indices as measured by the mouse grimace scale. This dissociation supports a model in which DAT⁺ vagal afferents preferentially regulate ongoing and affective–motivational dimensions of pain, whereas broader TH⁺ afferent populations exert more limited control over mechanical hypersensitivity (*39*). Together, these findings support a hierarchical and functionally segregated organization of dopaminergic vagal afferents in the modulation of trigeminal pain. Although recent single-cell transcriptomic studies of the nodose ganglion have provided valuable molecular classifications of vagal sensory neurons, dopamine-related transcripts such as Slc6a3 (DAT) have not consistently appeared as defining markers in some datasets. It is important to interpret this cautiously. Single-cell RNA sequencing can underestimate transcripts expressed at low levels or in relatively small neuronal populations, particularly when sequencing depth or clustering thresholds limit detection. As a result, the absence or low representation of a given transcript in a dataset does not necessarily indicate true biological absence.

Our decision to target DAT-expressing neurons was therefore guided not solely by transcriptomic prominence, but by molecular specificity. DAT encodes the dopamine transporter and is closely linked to regulated dopamine handling at synaptic sites. In contrast, broader TH expression may encompass neurons that do not uniformly support functional dopaminergic signaling (*40, 41*). Using a Cre-dependent strategy, we were able to selectively access this population. Reporter expression was confined to DAT-Cre–positive vagal afferents, and mCherry labeling confirmed the presence of a discrete DAT-positive neuronal subset within the nodose ganglion (Fig. 2). Importantly, chemogenetic activation of these neurons produced robust suppression of trigeminal nociceptive responses. These results position DAT-expressing vagal afferents as a functionally specialized component of the vagal sensory repertoire capable of modulating trigeminal pain circuits.

Whether vagal afferent nerves directly innervate the TMJ and convey TMJ-derived sensory information remains unexplored. Interestingly, an evolutionary-developmental perspective suggests that the auricular vagus nerve may have evolved from mandibular and jaw-associated sensory pathways (*22, 42*), raising the possibility that the vagus nerve plays a more fundamental role in TMJ function alongside trigeminal sensory nerves. Consistent with this view, we found that vagal afferents both anatomically and functionally innervate the TMJ and are activated by TMJ-related injury. Therefore, our findings that aVNS alleviates TMD-related pain support the notion that aVNS may be particularly effective for pain arising from the TMJ. Moreover, our data provide mechanistic insight into these analgesic effects, suggesting that involvement of dopaminergic vagal afferents contribute to both sensory and affective components of pain relief. Although the mechanisms underlying how the activation of dopaminergic vagal afferents suppresses pain signaling remains to be investigated, our data suggest a peripheral mechanism. Dopamine released from vagal afferents likely inhibits trigeminal afferent terminals in TMJ that are sensitized after injury. We believe this is a novel mechanism of analgesia exerted by vagus nerve stimulation that is distinct from, and in addition to, central mechanisms (*18*).

Several limitations should be acknowledged. Although our genetic access strategy targets vagal afferents with dopaminergic molecular features, it remains unclear, first, whether these neurons release dopamine at the TMJ, and second, which dopamine receptor subtypes mediate downstream effects. It is possible that additional co-transmitters or neuromodulatory mechanisms act in concert with dopaminergic signaling. Moreover, while our data establish a functional link between dopaminergic vagal afferent activation and modulation of trigeminal nociceptive processing, the precise anatomical locus and synaptic intermediates underlying this interaction remain unresolved, including whether vagal dopamine signaling acts directly at the level of the TMJ and TG or indirectly through brainstem circuits such as the nucleus tractus solitarius and associated neuromodulatory nuclei. Finally, translational work in humans combining vagal neuromodulation with neuroimaging may clarify whether analogous dopaminergic interoceptive mechanisms operate in clinical pain conditions. Altogether, our findings identify a distinct subset of vagal sensory afferents with dopaminergic molecular features that modulates trigeminal nociceptive processing and confers motivational significance to pain relief. These results provide a neurobiological basis for how internal physiological signals influence both sensory and affective dimensions of craniofacial pain and support the development of more precise, vagus-based neuromodulatory strategies.

## Acknowledgments

The authors thank Ms. Sinu Kumari for excellent technical assistance.

## Author contributions

Conceptualization: HS, DH, TL, MKC, YSK

Methodology: HS, DH, TL

Investigation: HS, DH, TL, CZ, MSSA, NB, JS, EK

Data curation: HS, DH, TL

Formal analysis: HS, DH, TL

Supervision: MKC, YSK

Funding acquisition: MKC, YSK

Writing – original draft: HS, DH, TL

Writing – review & editing: HS, DH, TL, MKC, YSK

## Data, code, and materials availability

The data that support the findings of this study are available from the corresponding authors upon reasonable request.

## Supplementary Materials

Materials and Methods

Supplementary Text

Figs. S1 to S2

Movies S1 to S12

## Materials and Methods

### Animals

All experimental procedures were approved by the Institutional Animal Care and Use Committee (IACUC) of the University of Alabama at Birmingham (UAB) and were performed in accordance with NIH guidelines for the care and use of laboratory animals. Adult male and female mice (8–12 weeks old) were used, including DAT-Cre (Slc6a3-Cre; The Jackson Laboratory, stock #006660) (*43*), TH-Cre (The Jackson Laboratory, stock #025614) (*44*), TrkC-Cre (Ntrk3-Cre; The Jackson Laboratory, stock #030291) (*45*), MrgD-Cre (Mrgprd-Cre; The Jackson Laboratory, stock #031286) (*46*), CGRP-Cre (from Pao-Tien Chuang, University of California, San Francisco) (*47*), and wild-type C57BL/6J (The Jackson Laboratory, stock #000664) (*48*). Animals were housed under standard laboratory conditions (12 h light/dark cycle, 22 ± 2°C) with ad libitum access to food and water.

### AAV-mediated chemogenetic targeting of vagal neurons

For selective activation of dopaminergic vagal neurons, AAV-DIO-hM3Dq (#44361; Addgene) or AAV-DIO-mCherry (control) (#50459; Addgene) was injected unilaterally into the right nodose ganglion of Cre-driver mice. Mice were anesthetized with isoflurane (2–3%) and placed in a stereotaxic frame. A small incision was made at the ventral neck, and the nodose ganglion was exposed under a surgical microscope. Viral vectors (0.5 μl administered on the right side) were injected using a pulled glass micropipette connected to a microinjection pump (Harvard Apparatus). Tamoxifen (75 mg/kg, 13258; Cayman Chemical) was administered once daily for five consecutive days, starting one week after viral injection, to induce Cre-dependent recombination. Mice were allowed to recover for three weeks after completion of tamoxifen treatment before behavioral experiments.

### Forced mouth opening (FMO) model

To induce TMJ injury and chronic orofacial pain, mice were anesthetized with an intraperitoneal injection of ketamine/xylazine (90/13.5 mg/kg; Zoetis, KET-11700002R2; VetOne, 33197) or isoflurane (3% induction, 1.5% maintenance) inhalation and subjected to FMO using a Colibri retractor (Fine Science Tools, 17000-03). The mouth was mechanically held open for 3 hours per day over 2 or 5 consecutive days to induce TMJ overextension without tissue damage. Control mice were anesthetized for the same duration but were not subjected to mouth opening.

### Chemogenetic activation

Chemogenetic activation of hM3Dq-expressing vagal neurons was achieved by intraperitoneal injection of compound 21 (C21, 0.5 mg/kg, HB6124; Hello Bio) according to the experimental schedule. C21 was administered at 5 p.m. on days 1, 2, 5, and 6 following the completion of the final FMO session, for a total of four injections.

### Transcutaneous auricular vagus nerve stimulation (aVNS)

aVNS was delivered under 2–3% isoflurane anesthesia. To prevent any mechanical impingement of ear skin, we used a concentric electrode (#BIOC-25; The Electrode Store), whose tip is attached to the ear with a conductive electrical pad (6494; XiBany). The pad was placed on the left cymba and cavum conchae—the territory of the auricular branch of the VN. A conductive gel (TensXtends; Flextone) was applied between the ear skin and the pad to ensure an electrical connection. Electrical stimuli were delivered through a TENS EM49 nerve stimulator (Beurer GmbH). The stimulator was connected to the concentric electrode through a concentric cable (BIOC-NH; The Electrode Store). The electrical stimulation parameter (frequency and intensity) was modified from our previous study (*31*). The daily aVNS was composed of two 20 min sessions in 3 hours apart. Each 20 min session was composed of electrical stimulation bursts (15 sec duration) applied at one-minute intervals. Each burst was composed of biphasic 2 mA pulses (250 us duration) at 2 Hz. In control-stimulated animals, the electrode remained attached to the conchae under isoflurane for the same period, but current was not delivered. aVNS and behavioral assays were conducted by two different investigators who were blinded to the experimental groups.

### TMJ von Frey test

The von Frey assay in the aVNS experiment was conducted as previously described (*49*). Animals were placed on a regular leather work glove worn by the experimenter and habituated for 20 minutes per day over 3–4 consecutive days. A series of calibrated von Frey filaments (0.008–4 g) was applied to the orofacial skin. Each filament was applied to the TMJ region five times at intervals of a few seconds. The force that elicited a 50% response frequency was subsequently calculated. For chemogenetic activation experiments using compound 21 (C21), mice were acclimated to the experimenter’s scent and gentle handling for two consecutive days prior to behavioral testing. Subsequently, they were habituated to the testing apparatus, consisting of a transparent Plexiglas chamber containing a 4-oz paper cup, for 2 hours per day over 3–5 consecutive days to minimize stress-induced variability. Baseline facial mechanical sensitivity was assessed once daily for 5–7 days using a series of von Frey filaments (0.04–1.4 g; Aesthesio) applied to the temporomandibular (TMJ) region. When the 50% withdrawal thresholds stabilized within the range of 0.5–0.7 g, this was designated as day 0. After induction of TMJ pain through five consecutive days of forced mouth opening (FMO), mechanical thresholds were re-assessed on days 1, 2, and 3 post-FMO using the up–down method. All assessments were performed at the same time each day to minimize circadian effects.

### Mouse Grimace Scale (MGS)

The Mouse Grimace Scale (MGS) assay for aVNS treatment was performed as previously described (*29*). Mice were placed individually in cubicles (5 × 5 × 5 cm) for 30 minutes, after which baseline measurements were recorded. Two digital video cameras (Sony HDR-CX230/B high-definition Handycam camcorders; Sony Corp., Tokyo, Japan) were positioned to maximize the likelihood of capturing clear facial images. At each experimental time point, mice were videotaped for 30 minutes. During video recording, a clear image of the entire face was manually captured every 2–3 minutes (10 images per 30-minute session) using a still camera. For chemogenetic activation experiments using compound 21 (C21), mice were placed individually in a 4-oz paper cup and allowed to acclimate for 15 minutes before image acquisition. Facial images were captured using a high-resolution digital camera placed at a fixed distance and angle throughout the experiment to ensure consistency. Images were taken at baseline and on days 5 and 7 after FMO. Five facial action units—orbital tightening, ear position, nose bulge, cheek bulge, and whisker change—were scored. Each action unit was rated on a scale of 0, 1, or 2 according to established criteria by experimenters blinded to the treatment conditions.

### Open field test (OFT)

Open field testing was conducted to evaluate locomotor activity and anxiety-like behaviors following FMO. Mice were brought into the testing room and allowed to acclimate for at least 30 minutes before testing. The open-field arena measured 40 × 40 × 30 cm and was constructed with opaque acrylic walls. Each mouse was gently placed in the center of the arena to begin the session. Testing was performed during the light phase (08:00–14:00) to minimize circadian variation. Each trial lasted 10 minutes and was recorded using a top-mounted video camera, and behavioral tracking and quantitative analysis were carried out with ToxTrac software (*50*). The arena was cleaned with 70% ethanol between sessions to eliminate odor cues. Measurements were performed at baseline and on days 5 and 7 after FMO to assess changes in exploratory and anxiety-like behaviors.

### Conditioned Place Preference (CPP)

The conditioned place preference test was performed to assess motivational valence associated with dopaminergic vagal activation. The CPP apparatus consisted of two distinct chambers (chamber A and B) with different visual and tactile cues separated by a removable partition. On day 1 (pre-test), mice were allowed to freely explore both chambers for 15 minutes to establish baseline preference. During the conditioning phase (days 2–4), in the morning, mice were placed in chamber A and received saline injections (i.p. or intra-TMJ), and in the afternoon, mice were placed in chamber B and received C21 (0.5 mg/kg, i.p., or intra-TMJ), with a 6-hour interval between sessions. Each session lasted 15 minutes, and mice were confined to the corresponding chamber immediately after injection. On day 5 (test phase), mice were again allowed free access to both chambers for 15 minutes, and their activity was recorded using a top-mounted camera. CPP scores were calculated as the change in time spent in the C21-paired chamber between the pre-test and test sessions (ΔCPP = post – pre). All experiments were conducted under consistent lighting and noise conditions, and chambers were cleaned with 70% ethanol between trials. A positive ΔCPP value indicated a rewarding effect of dopaminergic vagal activation, whereas a negative value reflected aversion.

### In vivo Pirt-GCaMP3 Ca2+ imaging of intact trigeminal ganglia (TG)

In vivo calcium imaging of the intact TG was conducted in DAT-Cre/Pirt-GCaMP3 mice, following procedures adapted from previous work (*30, 51–54*). After surgical exposure of the TG, mice were positioned on a custom-built imaging platform and maintained at 37 ± 0.5°C using a heating pad. Anesthesia was maintained with 1–2% isoflurane in pure oxygen via a vaporizer. Live calcium signals were recorded for approximately 2–3 hours using a confocal microscope equipped with solid-state diode lasers (excitation = 488 nm, emission = 500–550 nm). Image stacks were acquired at 512 × 512 pixels or higher resolution using a 5×/0.25 NA dry objective, with frame rates of 4.5–8.8 s per frame, across depths of 0–900 µm. Mechanical stimuli (0.4 g and 2 g von Frey filaments and a small brush), thermal stimuli (noxious water at 4°C and 50°C), and chemical stimulation (10 µL of 500 µM capsaicin injected intracutaneously) were applied to the jaw region. Raw imaging sequences were deconvolved and realigned using the StackReg plugin in ImageJ (NIH), employing rigid-body cross-correlation for motion correction. Calcium transients were quantified as ΔF/F₀, ΔF equals Ft-F0 where Fₜ is the fluorescence at each frame and F₀ is the baseline fluorescence averaged across the first four frames. Responsive neurons were visually verified from the processed image stacks.

### Retrograde labeling of TMJ afferents and immunohistochemistry

Wheat germ agglutinin (WGA) conjugated with Alexa Fluor 488 (WGA-488; Thermo-Fisher Scientific, Invitrogen, Carlsbad, CA) was injected into the TMJ to retrogradely label TMJ afferents. Procedurally, fur was shaved in the TMJ area, and the location of the TMJ was identified by following the posterior ramus of the mandible and the end of the zygomatic arch. Under ketamine/xylazine anesthesia, WGA-488 (1% in distilled water; 4 µl/joint) was injected into the joint cavity of the TMJ. The mice were euthanized 3 days after the injection. Trigeminal ganglia and vagal ganglia were collected and cryosectioned, and immunohistochemical assays were performed using the antibody against CGRP.

### Tissue processing and confocal fluorescence imaging

Mice were deeply anesthetized and transcardial perfusion was performed with cold phosphate-buffered saline (PBS), followed by 4% paraformaldehyde (PFA). The nodose ganglia (NG) were carefully dissected, post-fixed in 4% PFA at 4°C for 24 hours and cryoprotected in 30% sucrose solution overnight. Tissues were embedded in OCT compound and stored at -80°C until sectioning. Serial sections of 15 µm thickness were prepared using a cryostat (Leica) and mounted onto glass slides. For immunofluorescence staining, sections were washed with PBS and permeabilized in PBST (0.3% Triton X-100 in PBS). Samples were then blocked with 5% normal goat serum for 1 hour at room temperature. mCherry fluorescence, expressed in a Cre-dependent manner following viral infection, was detected directly without antibody-based amplification. Nuclei were counterstained with DAPI ready-made solution (MBD0015; Millipore Sigma). Sections were rinsed in PBS and coverslipped with ProLong™ Diamond Antifade Mountant (Invitrogen). Images were acquired using a confocal microscope (Zeiss LSM 880) equipped with 10× dry and 40× water-immersion objectives, and all parameters (laser power, gain, offset) were kept constant across samples.

### Statistical analysis

All data are expressed as mean ± s.e.m. Statistical analyses were performed using GraphPad Prism 10. Behavioral data were analyzed using two-way ANOVA with repeated measures, followed by Tukey’s post hoc test. MGS and CPP data were analyzed by unpaired t-tests or one-way ANOVA were appropriate. P < 0.05 was considered statistically significant.

---

**Fig. S1.**
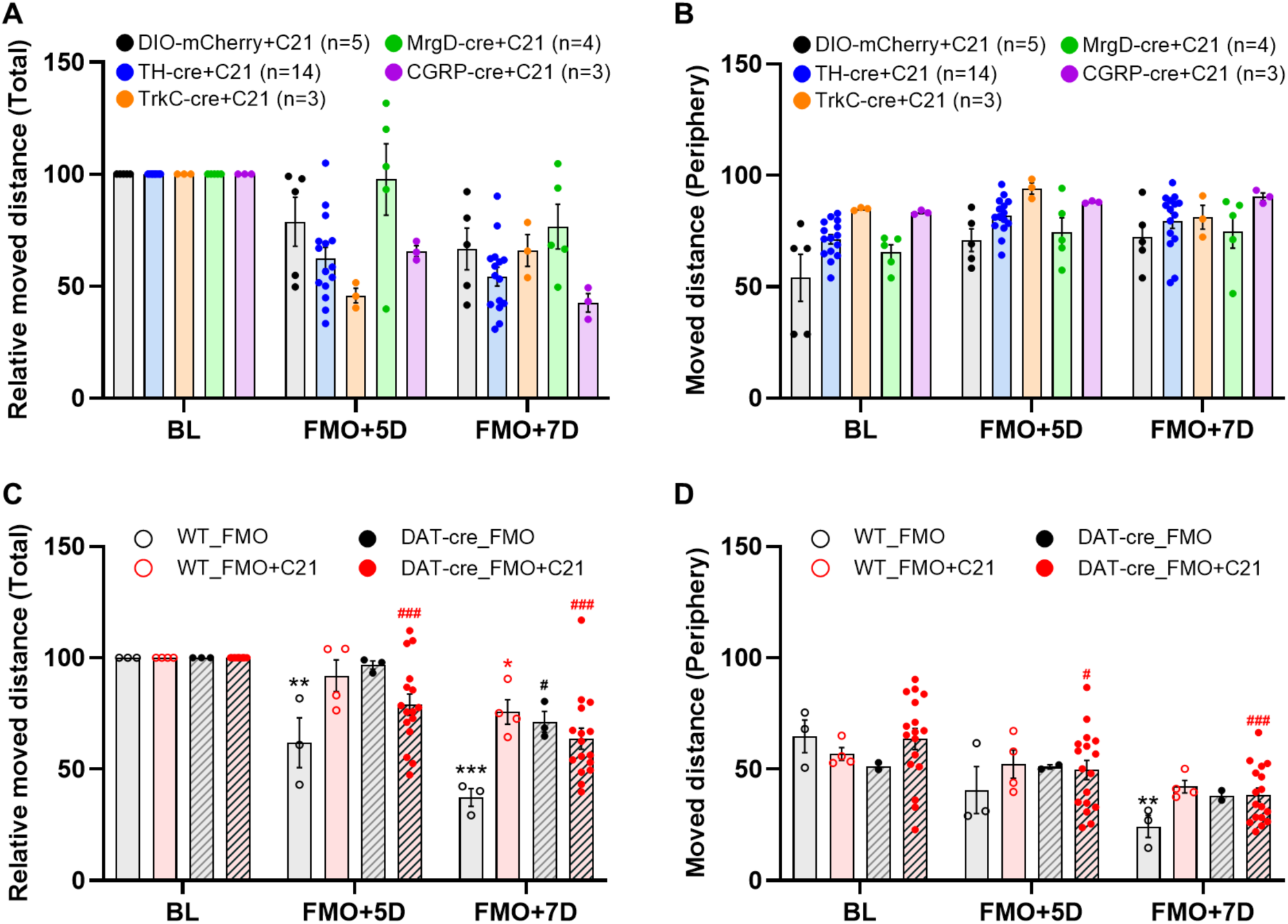
Chemogenetic activation does not alter locomotor activity after FMO. (A, **B)** Locomotor activity measured by total distance and peripheral distance in different Cre lines after FMO. No major changes were observed across genotypes. **(C and D)** Comparison of locomotor indices between WT and DAT-Cre mice with or without C21 treatment confirmed that chemogenetic activation did not impair motor function. Data are mean ± s.e.m.; *p < 0.05, **p < 0.01, ***p < 0.001, #p < 0.05, ###p < 0.001 (two-way ANOVA followed by Tukey’s multiple-comparisons test).

**Fig. S2.**
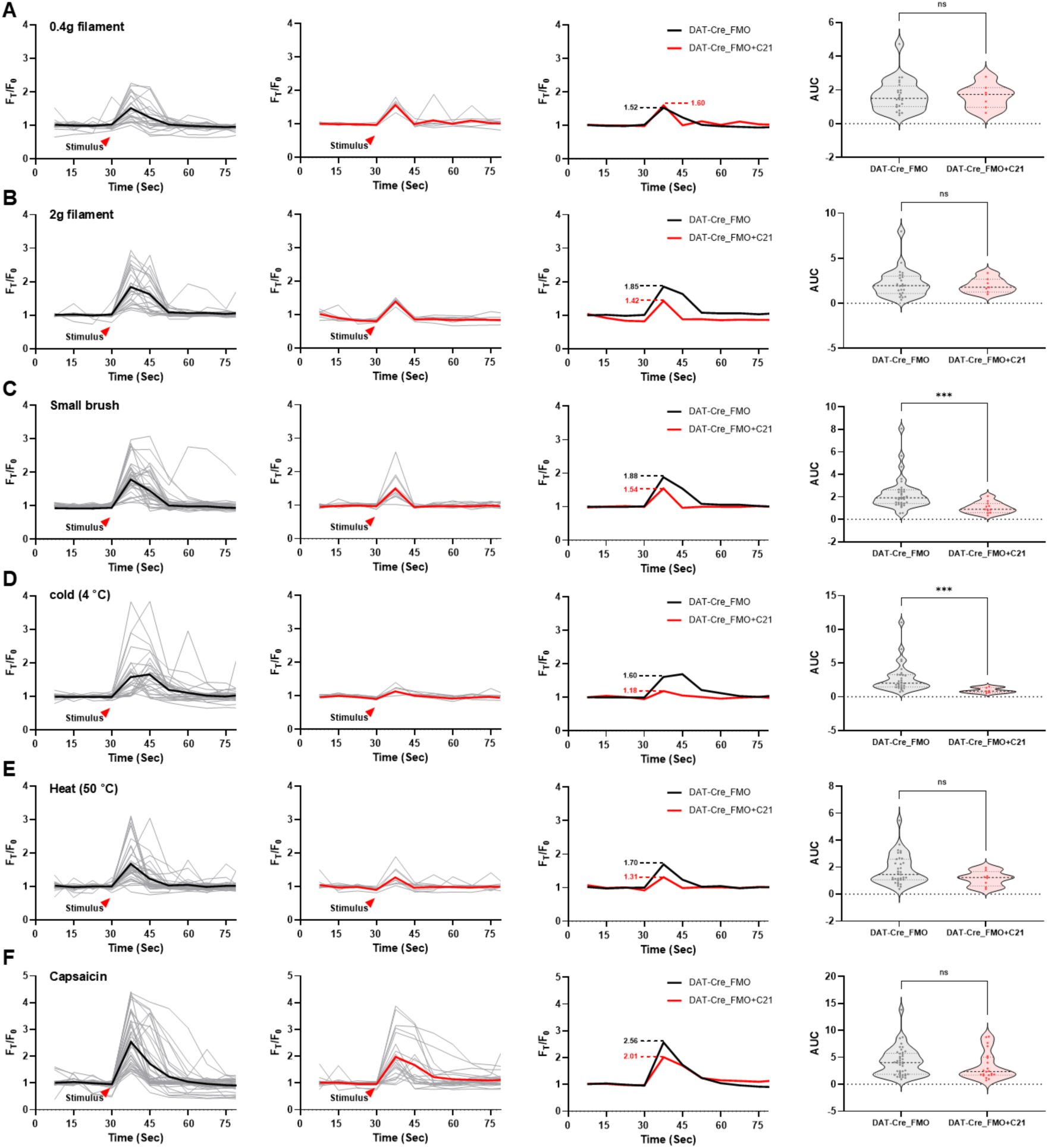
Superimposed normalized TG Ca²⁺ transients evoked by V3 stimuli after FMO with or without C21 treatment. (A–F) Representative superimposed normalized Ca²⁺ transient traces recorded from TG neurons in DAT-Cre/GCaMP3 mice subjected to FMO, with or without C21 treatment (DREAD virus was injected), in response to different V3 stimuli: **(A)** 0.4 g von Frey filament, **(B)** 2 g von Frey filament, **(C)** small brush, **(D)** cold (4 °C), **(E)** heat (50 °C), and **(F)** capsaicin. Thin grey lines indicate individual neuronal responses, and bold lines represent the averaged Ca²⁺ transients for each group (black without C21 or red with C21. Red arrowheads denote the timing of stimulus application. Right panels show violin plots summarizing the area under the curve (AUC) of normalized Ca²⁺ transients for each stimulus condition. Dotted lines indicate quartiles and the central dotted line indicates the median. Statistical analyses were performed using the Mann–Whitney test; ***p < 0.001; ns, not significant.

**Movie S1, related to Figure. 4D**

Representative in vivo GCaMP Ca^2+^ imaging of intact Trigeminal Ganglion (TG) of spontaneous activity (no stimulus) using DAT-Cre+Forced Mouth Opening (FMO) mouse after DREAD virus injection. Note that the resolution of the movie was reduced by conversion from Zen blue (Zeiss) software to MP4 movie format.

**Movie S2, related to Figure. 4D**

Representative in vivo GCaMP Ca2+ imaging of intact TG of spontaneous activity (no stimulus) using DAT-Cre+FMO+C21 mouse.

**Movie S3, related to Figure. 4E**

Representative in vivo GCaMP Ca2+ imaging of intact TG of activated neurons in response to 0.4 g filament stimulus at V3 region using DAT-Cre+FMO mouse.

**Movie S4, related to Figure. 4E**

Representative in vivo GCaMP Ca2+ imaging of intact TG of activated neurons in response to 0.4 g filament stimulus at V3 region using DAT-Cre+FMO+C21 mouse.

**Movie S5, related to Figure. 4F**

Representative in vivo GCaMP Ca2+ imaging of intact TG of activated neurons in response to 2 g filament stimulus at V3 region using DAT-Cre+FMO mouse.

**Movie S6, related to Figure. 4F**

Representative in vivo GCaMP Ca2+ imaging of intact TG of activated neurons in response to 2 g filament stimulus at V3 region using DAT-Cre+FMO+C21 mouse.

**Movie S7, related to Figure. 4G**

Representative in vivo GCaMP Ca^2+^ imaging of intact TG of activated neurons in response to small brush stimulus at V3 region using DAT-Cre+FMO mouse.

**Movie S8, related to Figure. 4 G**

Representative in vivo GCaMP Ca^2+^ imaging of intact TG of activated neurons in response to small brush stimulus at V3 region using DAT-Cre+FMO+C21 mouse.

**Movie S9, related to Figure. 4I**

Representative in vivo GCaMP Ca^2+^ imaging of intact TG of activated neurons in response to Heat (50 °C water) stimulus at V3 region using DAT-Cre+FMO mouse.

**Movie S11, related to Figure. 4J**

Representative in vivo GCaMP Ca^2+^ imaging of intact TG of activated neurons in response to capsaicin at V3 region using DAT-Cre+FMO mouse.

**Movie S12, related to Figure. 4J**

Representative in vivo GCaMP Ca^2+^ imaging of intact TG of activated neurons in response to capsaicin at V3 region using DAT-Cre+FMO+C21 mouse.

## References and Notes

1. A. D. Craig, Interoception: The sense of the physiological condition of the body. Curr. Opin. Neurobiol. 13, 500–505 (2003). doi: 10.1016/s0959-4388(03)00090-4; pmid: 12965300

2. R. L. Wang, R. B. Chang, The coding logic of interoception. Annu. Rev. Physiol. 86, 301–327 (2024). doi: 10.1146/annurev-physiol-042222-023455; pmid: 38061018

3. M. M. Kaelberer et al., A gut-brain neural circuit for nutrient sensory transduction. Science 361, eaat5236 (2018). doi: 10.1126/science.aat5236; pmid: 30237325

4. H.-E. Tan et al., The gut-brain axis mediates sugar preference. Nature 580, 511–516 (2020). doi: 10.1038/s41586-020-2199-7; pmid: 32322067

5. M. Li et al., Gut-brain circuits for fat preference. Nature 610, 722–730 (2022). doi: 10.1038/s41586-022-05266-z; pmid: 36070796

6. W. Han et al., A Neural Circuit for Gut-Induced Reward. Cell 175, 665–678.e23 (2018). doi: 10.1016/j.cell.2018.08.049; pmid: 30245012

7. H. Jin et al., A body–brain circuit that regulates body inflammatory responses. Nature 630, 695–703 (2024). doi: 10.1038/s41586-024-07469-y; pmid: 38692285

8. C. W. Austelle et al., Vagus nerve stimulation (VNS): recent advances and future directions. Clin. Auton Res 34, 529–547 (2024). doi: 10.1007/s10286-024-01065-w; pmid: 39363044

9. K. Chakravarthy et al., Review of the Uses of Vagal Nerve Stimulation in Chronic Pain Management. Curr Pain Headache Rep 19, 54 (2015). doi: 10.1007/s11916-015-0528-6; pmid: 26493698

10. V. Costa et al., Transcutaneous vagus nerve stimulation effects on chronic pain: systematic review and meta-analysis. Pain Rep 9, e1171 (2024). doi: 10.1097/PR9.0000000000001171; pmid: 39131814

11. I. Lerman et al., Noninvasive vagus nerve stimulation alters neural response and physiological autonomic tone to noxious thermal challenge. PLOS ONE 14, e0201212 (2019). doi: 10.1371/journal.pone.0201212; pmid: 30759089

12. P. Shao et al., Role of Vagus Nerve Stimulation in the Treatment of Chronic Pain. Neuroimmunomodulation 30, 167–183 (2023). doi: 10.1159/000531626; pmid: 37369181

13. Y. Guo, P. Gharibani, Analgesic Effects of Vagus Nerve Stimulation on Visceral Hypersensitivity: A Direct Comparison Between Invasive and Noninvasive Methods in Rats. Neuromodulation 27, 284–294 (2024). doi: 10.1016/j.neurom.2023.04.001; pmid: 37191611

14. B. E. Cairns, Pathophysiology of TMD pain - basic mechanisms and their implications for pharmacotherapy. J. Oral Rehabil 37, 391–410 (2010). doi: 10.1111/j.1365-2842.2010.02074.x; pmid: 20337865

15. F. P. Kapos et al., Temporomandibular disorders: a review of current concepts in aetiology, diagnosis and management. Oral Surg 13, 321–334 (2020). doi: 10.1111/ors.12473; pmid: 34853604

16. A. Herrero Babiloni et al., Temporomandibular disorders cases with high-impact pain are more likely to experience short-term pain fluctuations. Sci Rep 12, 1657 (2022). doi: 10.1038/s41598-022-05598-w; pmid: 35102207

17. A. U. Yap et al., Characteristics of painful temporomandibular disorders and their influence on jaw functional limitation and oral health-related quality of life. J Oral Rehabil 51, 1748–1758 (2024). doi: 10.1111/joor.13768; pmid: 38845181

18. L. E. Cornelison et al., Inhibition of Trigeminal Nociception by Non-invasive Vagus Nerve Stimulation: Investigating the Role of GABAergic and Serotonergic Pathways in a Model of Episodic Migraine. Front Neurol 11, 593612 (2020). doi: 10.3389/fneur.2020.00146; pmid: 32194498

19. J. L. Hawkins et al., Vagus nerve stimulation inhibits trigeminal nociception in a rodent model of episodic migraine. Pain Rep 2, e628 (2017). doi: 10.1097/PR9.0000000000000628: pmid: 29392242

20. Y. Huang et al. The modulation effects of repeated transcutaneous auricular vagus nerve stimulation on the functional connectivity of key brainstem regions along the vagus nerve pathway in migraine patients. Front Mol Neurosci 16, 1160006 (2023). doi: 10.3389/fnmol.2023.1160006; pmid: 37333617

21. J. Y. Y. Yap et al., Critical Review of Transcutaneous Vagus Nerve Stimulation: Challenges for Translation to Clinical Practice. Front Neurosci 14, 284 (2020). doi: 10.3389/fnins.2020.00284; pmid: 32410932

22. E. Kaniusas et al., Current Directions in the Auricular Vagus Nerve Stimulation I - A Physiological Perspective. Front Neurosci 13, 854 (2019). doi: 10.3389/fnins.2019.00854; pmid: 31447643

23. L. S. Prott et al., Transcutaneous auricular vagus nerve stimulation for the treatment of myoarthropatic symptoms in patients with craniomandibular dysfunction – a protocol for a randomized and controlled pilot trial. Pilot Feasibility Stud 10, 27 (2024). doi: 10.1186/s40814-024-01447-x; pmid: 38331976

24. A. Percin et al., The effect of auricular vagus nerve stimulation in women with temporomandibular joint disorders: a randomized controlled study. Rev Assoc Med Bras (1992) 71, e20241739 (2025). doi: 10.1590/1806-9282.20241739; pmid: 40465996

25. Y.S. Kim et al., Coupled Activation of Primary Sensory Neurons Contributes to Chronic Pain. Neuron 91, 1085–1096 (2016). doi: 10.1016/j.neuron.2016.07.044; pmid: 27568517

26. Y. Zhang et al., Imaging sensory transmission and neuronal plasticity in primary sensory neurons with a positively tuned voltage indicator. Nat Commun 16, 6396 (2025). doi: 10.1038/s41467-025-61774-2; pmid: 40640176

27. J. Shannonhouse et al., In Vivo Calcium Imaging of Neuronal Ensembles in Networks of Primary Sensory Neurons in Intact Dorsal Root Ganglia. J Vis Exp (192) (2023). doi: 10.3791/64826; pmid: 36847407

28. J. Shannonhouse et al., Lessons from the use of in vivo cellular calcium imaging in primary sensory neurons and spinal cord. Neuroscientist 31, 591–610 (2025). doi: 10.1177/10738584251360724; pmid: 40804523

29. I. Alshanqiti et al., Posttraumatic hyperalgesia and associated peripheral sensitization after temporomandibular joint injury in mice. Pain 166, 1597 (2025). doi: 10.1097/j.pain.0000000000003498; pmid: 39715145

30. H. Son et al., Elucidation of neuronal activity in mouse models of temporomandibular joint injury and inflammation by in vivo GCaMP Ca^2+^ imaging of intact trigeminal ganglion neurons. Pain 165, 2794–2803 (2024). doi: 10.1097/j.pain.0000000000003421; pmid: 39365648

31. M. S. S. Ali et al., Genetic labeling of the nucleus of tractus solitarius neurons associated with electrical stimulation of the cervical or auricular vagus nerve in mice. Brain Stimul 17, 987–1000 (2024). doi: 10.1016/j.brs.2024.08.007; pmid: 39173736

32. W. He, et al., Auricular Acupuncture and Vagal Regulation. Evid Based Complement Alternat Med 2012, 786839 (2012). doi: 10.1155/2012/786839; pmid: 23304215

33. J. Kupari et al., An Atlas of Vagal Sensory Neurons and Their Molecular Specialization. Cell Rep 27, 2508–2523.e4 (2019). doi: 10.1016/j.celrep.2019.04.096; pmid: 31116992

34. L. Bai et al., Genetic Identification of Vagal Sensory Neurons That Control Feeding. Cell 179, 1129–1143.e23 (2019). doi: 10.1016/j.cell.2019.10.031; pmid: 31730854

35. E. Kim et al., BoNT Injection into Temporomandibular Joint Alleviates TMJ Pain in Forced Mouth Opening Mouse Model. J Neurosci 45, e2035242025 (2025). doi: 10.1523/JNEUROSCI.2035-24.2025; pmid: 40759513

36. B. Rubí, P. Maechler, Minireview: New Roles for Peripheral Dopamine on Metabolic Control and Tumor Growth: Let’s Seek the Balance. Endocrinology 151, 5570–5581 (2010). doi: 10.1210/en.2010-0745; pmid: 21047943

37. A. Khaky et al., Tyrosine Hydroxylase-Expressing Neurons in the Vagal Ganglia: Characterization and Implications. Biomedicines 13, 2126 (2025). doi: 10.3390/biomedicines13092126; pmid: 41007689

38. M. N. Baliki, A. V. Apkarian, Nociception, pain, negative moods, and behavior selection. Neuron 87, 474–491 (2015). doi: 10.1016/j.neuron.2015.06.005; pmid: 26247858

39. E. Navratilova, F. Porreca, Reward and motivation in pain and pain relief. Nat Neurosci 17, 1304–1312 (2014). doi: 10.1038/nn.3811; pmid: 25254980

40. G. E. Torres, R. R. Gainetdinov, M. G. Caron, Plasma membrane monoamine transporters: structure, regulation and function. Nat Rev Neurosci 4, 13–25 (2003). doi: 10.1038/nrn1008; pmid: 12511858

41. R. A. Vaughan, J. D. Foster, Mechanisms of dopamine transporter regulation in normal and disease states. Trends Pharmacol Sci 34, 489–496 (2013). doi: 10.1016/j.tips.2013.07.005; pmid: 23968642

42. J. Meng, Y. Wang, C. Li, Transitional mammalian middle ear from a new Cretaceous Jehol eutriconodont. Nature 472, 181–185 (2011). doi: 10.1038/nature09921; pmid: 21490668

43. C. M. Bäckman et al., Characterization of a mouse strain expressing Cre recombinase from the 3′ untranslated region of the dopamine transporter locus. Genesis 44, 383–390 (2006). doi: 10.1002/dvg.20228; pmid: 16865686

44. V. E. Abraira et al., The cellular and synaptic architecture of the mechanosensory dorsal horn. Cell 168, 295–310.e19 (2017). doi: 10.1016/j.cell.2016.12.010; pmid: 28041852

45. L. Bai et al., Genetic identification of an expansive mechanoreceptor sensitive to skin stroking. Cell 163, 1783–1795 (2015). doi: 10.1016/j.cell.2015.11.060; pmid: 26687362

46. Olson, W. et al. Sparse genetic tracing reveals regionally specific functional organization of mammalian nociceptors. Elife 6, e29507 (2017). doi: 10.7554/eLife.29507; pmid: 29022879

47. H. Song et al., Functional characterization of pulmonary neuroendocrine cells in lung development, injury, and tumorigenesis. Proc Natl Acad Sci U S A 109, 17531–17536 (2012). doi: 10.1073/pnas.1207238109; pmid: 23047698

48. B. Paigen et al., Variation in susceptibility to atherosclerosis among inbred strains of mice. Atherosclerosis 57, 65–73 (1985). doi: 10.1016/0021-9150(85)90138-8; pmid: 3841001

49. S. Wang et al., Ablation of TRPV1⁺ afferent terminals by capsaicin mediates long-lasting analgesia for trigeminal neuropathic pain. eNeuro 7, ENEURO.0118-20.2020 (2020). doi: 10.1523/ENEURO.0118-20.2020; pmid: 32404326

50. A. Rodriguez et al., ToxTrac: A fast and robust software for tracking organisms. Methods Ecol Evol 9, 460–464 (2018). doi: 10.1111/2041-210x.12874

51. J. Shannonhouse et al., In Vivo Calcium Imaging of Neuronal Ensembles in Networks of Primary Sensory Neurons in Intact Trigeminal Ganglia. J Vis Exp (222) (2025). doi: 10.3791/68284; pmid: 40824880

52. Y.S. Kim et al., Central terminal sensitization of TRPV1 by descending serotonergic facilitation modulates chronic pain. Neuron 81, 873–887 (2014). doi: 10.1016/j.neuron.2013.12.011; pmid: 24462040

53. H. Son et al., PACAP38/mast-cell-specific receptor axis mediates repetitive stress-induced headache in mice. J Headache Pain 25, 87 (2024). doi: 10.1186/s10194-024-01786-3; pmid: 38802819

54. Y. Zhang et al., In vivo Pirt-Marina voltage sensor imaging detects primary sensory neuron-specific voltage dynamics and neuronal plasticity changes. Proc Natl Acad Sci U S A 122, e2416712122 (2025). doi: 10.1073/pnas.2416712122; pmid: 40938705

